# Predicting Lifespan-Extending Compounds Using Supervised Machine Learning

**DOI:** 10.64898/2026.10.16.759743

**Authors:** Divyam Acharya, Alex A. Freitas

## Abstract

Identifying lifespan-extending compounds is an important objective in ageing research, but experimental screening remains costly and time-consuming. This study evaluates supervised machine learning for predicting lifespan-extending compounds in *Caenorhabditis elegans* using compound–protein interaction features derived from the STITCH database. Three feature representations were compared: continuous interaction probabilities, binary features generated using a fixed threshold of 0.5, and binary features generated using an optimised threshold selected within cross-validation. Random Forest and XGBoost classifiers were evaluated using stratified 10-fold cross-validation, with performance assessed by accuracy, area under the receiver operating characteristic curve (AUROC), and area under the precision– recall curve (AUPRC). Model interpretation was examined using Mean Decrease in Impurity (MDI), SHAP, TreeSHAP, and LIME, with Jaccard similarity used to compare the top-ranked protein features. Random Forest achieved its highest accuracy (78.84%), AUROC (0.7391), and AUPRC (0.5202) using the fixed-threshold binary representation. XGBoost achieved its highest accuracy (80.36%) using the optimised-threshold representation, whereas continuous features yielded its highest AUROC (0.7605) and AUPRC (0.5332). SHAP and TreeSHAP showed the strongest agreement in protein rankings, producing identical top-10 feature sets across all three XGBoost representations. These findings demonstrate that feature representation influences both predictive performance and model interpretation, highlighting the importance of evaluating classification metrics alongside explainability when prioritising candidate compounds for further ageing research.

## 1. Introduction

Ageing is a complex biological process characterised by the gradual decline of physiological function and an increased risk of disease (Partridge, 2010). As global life expectancy continues to rise, ageing-related disorders are becoming increasingly prevalent. Consequently, identifying interventions capable of extending healthy lifespan has become a major goal of biomedical research.

One promising approach involves the discovery of pharmacological compounds that influence biological pathways associated with ageing. Experimental studies have identified compounds capable of extending lifespan in model organisms (Partridge, 2010; Chmielewski et al., 2025). However, experimentally evaluating large numbers of candidate compounds remains expensive, labour intensive, and time consuming. *Caenorhabditis elegans* has become one of the most widely used model organisms for ageing research due to its short lifespan, low experimental cost, and well characterised genetics (Ribeiro et al., 2023). Furthermore, many biological mechanisms associated with ageing are conserved between *C. elegans* and higher organisms, making it a useful model for investigating lifespan-extending interventions (Partridge, 2010).

By analysing data in biological databases, machine learning models can identify predictive patterns associated with lifespan extension and help prioritise compounds for future experimental investigation (Auslander, Gus-sow and Koonin, 2021). In particular, ensemble learning methods such as Random Forest and XGBoost have shown considerable success in biological prediction tasks due to their ability to model complex relationships within high-dimensional datasets while remaining relatively robust to overfitting. However, many machine learning studies in biology focus primarily on predictive accuracy while providing limited insight into the biological mechanisms underlying model predictions (Chen et al., 2024).

Interpreting classification models is especially important in biomedical applications because researchers require explanations that can guide future experimental investigation. Explainable Artificial Intelligence (XAI) methods provide mechanisms for understanding how machine learning models generate predictions and which biological features contribute most strongly to classification decisions (Chen et al., 2024). Such methods are particularly valuable in ageing research because they may help identify proteins and pathways potentially associated with lifespan regulation. This project addresses the supervised machine learning (classification) task of predicting whether or not a compound extends the lifespan of *C. elegans* using biologically interpretable protein–compound interaction features. In addition to evaluating predictive performance, the project places strong emphasis on explainability and biological interpretation through systematic feature importance analysis.

Three types of feature representations were investigated:

- Continuous probability-based protein-interaction features.
- Binary protein-interaction features generated using a fixed threshold of 0.5.
- Binary protein-interaction features generated using an optimised threshold selected during cross-validation.

This research has three objectives. The first objective is to compare the effect of these three different types of feature representations on the predictive performance of the classification models learned by two popular and strong types of classification algorithms: random forests and XGBoost.

The second objective is to identify the most important features in the learned models, using four explainability methods – more specifically, four feature im-portance methods described in Section 2.3; i.e., to identify the top-ranked proteins consistently associated with lifespan-extending compounds across the three above feature representations and two types of classification models (Random Forest and XGBoost).

The third objective, which is also related to explainability, is to analyse to what extent different XAI methods (feature importance measures) produce similar results in terms of identifying the most important features (proteins) in a given classification model.

Overall, by combining supervised machine learning with XAI techniques, this interdisciplinary project aims to contribute both computational and biological understanding to the study of ageing-related pharmacological interventions.

The remainder of this paper is structured as follows. Section 2 provides relevant background and related concepts. Section 3 describes the datasets, feature representations, machine learning models, evaluation metrics, and explainability methods used in this study. Section 4 presents the experimental results. Section 5 discusses the findings in relation to the research objectives. Section 6 concludes the report, while Section 7 outlines the limitations of the study and directions for future work.

## 2. Background

In recent years, machine learning methods have become increasingly important within biomedical research because of their ability to analyse complex biological datasets containing thousands of variables simultaneously. Biological systems are highly interconnected and often involve nonlinear interactions between genes, proteins, signalling pathways, and chemical compounds, making traditional statistical approaches insufficient for capturing these relationships effectively (Auslander, Gussow and Koonin, 2021). Consequently, supervised machine learning methods such as Random Forests and XGBoost have emerged as powerful tools for identifying hidden biological patterns and generating predictive models that can support ageing research.

### 2.1 Predicting whether or not a compound (drug) extends lifespan

There has been growing interest in applying machine learning techniques to the prediction of lifespan-extending compounds using datasets derived from DrugAge, a database containing information about compounds that extend lifespan in several model organisms, together with related biological databases. Barardo et al. (2017) demonstrated that Random Forest models can successfully predict lifespan-extending compounds in *C. elegans* using DrugAge data, achieving an accuracy of approximately 80%.

More recently, Kapsiani and Howlin (2021) further demonstrated the effectiveness of Random Forest for lifespan prediction by developing classifiers based on DrugAge compounds and molecular descriptors, achieving an AUC of 0.815 and successfully identifying candidate compounds with potential longevity-promoting effects. Their findings highlighted the suitability of machine learning methods for prioritising compounds for further experimental investigation. Ribeiro et al. (2023) extended this line of research by investigating biologically interpretable features derived from compound–protein interactions and Gene Ontology annotations, demonstrating that machine learning models can achieve strong predictive performance while also providing bio-logically meaningful explanations of their predictions.

This project was strongly inspired by the methodology proposed by Ribeiro et al. (2023), particularly the use of biologically interpretable compound–protein (drug-protein) interaction features rather than conventional chemical descriptors. However, that work used only one representation of compound– protein interactions, applying a fixed threshold (0.5 probability) to binarise continuous interaction probabilities. This project extends that research by investigating the effect of three alternative feature representations on both predictive performance and the identification of important proteins within the resulting classification models, as outlined in the Introduction.

### 2.2 Classification Algorithms: Random Forest and XG-Boost

Random Forest was one of the two machine learning algorithms used in this project because it is well suited to high-dimensional biological datasets and can model complex nonlinear relationships while remaining resistant to overfitting (Chen et al., 2024; Bulut et al., 2023). As an ensemble learning method, Random Forest constructs multiple decision trees using bootstrap sampling and random feature selection before aggregating predictions across the ensemble (Breiman, 2001). This approach reduces variance and improves predictive robustness compared with individual decision trees, making it particularly suitable for datasets containing large numbers of biological features. Furthermore, Random Forest naturally provides feature importance measures, allowing investigation of the proteins contributing most strongly to model predictions and making it particularly suitable for explainability analysis.

XGBoost (Extreme Gradient Boosting) was selected as the second algorithm to complement Random Forest through a different ensemble strategy. Unlike Random Forest which builds trees independently, XGBoost constructs trees sequentially using gradient boosting, where each new tree corrects the errors of previous ones (Chen and Guestrin, 2016). This sequential approach often captures more complex biological patterns. XGBoost offers several advantages for this study: regularisation to prevent overfitting on high dimensional data, native handling of sparse protein interaction probabilities, and built in feature importance measures. Both Random Forest and XGBoost enable comparison between bagging and boosting ensemble methods, providing a more comprehensive evaluation of predictive performance and identification of the most important features for lifespan-extending compounds.

### 2.3 eXplainable AI (XAI): Feature Importance Mea-sures

In addition to predictive modelling, a major focus of this project was explain-ability and biological interpretation. Although machine learning models may achieve strong predictive accuracy, understanding why predictions are made is equally important in biomedical research. Chen et al. (2024) emphasised that the explainability of machine learning models are crucial for building trust in their results within biological and medical domains, and they identified common pitfalls such as relying on a single explanation method or presenting cherry picked results. Consequently, this work investigates the use of four Explainable Artificial Intelligence (XAI) methods – more specifically, four feature importance methods – for identifying proteins (features) most strongly associated with lifespan-extending compounds. Each of these four methods first ranks the features in decreasing order of importance (according to a certain measure) in a given classification model, and then identifies the top-ranked features, which can then be shown to biologists in real-world applications (note, however, that a biological analysis of the top-ranked features is out of the scope of this project).

The first of these feature-importance methods was Mean Decrease in Impurity (MDI), which is the default feature importance measure embedded within Random Forest models. MDI estimates how much each feature contributes to reducing class impurity across all decision trees in the ensemble (Breiman, 2001), with features that consistently generate informative decision splits receiving higher importance scores. Despite known limitations, MDI remains one of the most widely used measures of feature importance due to its computational efficiency and natural integration within the Random Forest framework.

The second method was SHAP (SHapley Additive exPlanations), an explainability framework based on cooperative game theory that estimates the contribution of individual features to model predictions while satisfying desirable properties such as local accuracy and consistency (Lundberg and Lee, 2017). However, Shapley-value-based explanations have been criticised for several conceptual and mathematical limitations. Kumar et al. (2020) argued that feature attributions can vary depending on whether conditional or interventional distributions are used and noted that Shapley-value-based explanations do not naturally align with human-centred notions of explanation. More recently, Hwang et al. (2025) demonstrated that SHAP-based explanations are sensitive to feature representation, showing that common data engineering operations such as bucketisation and feature encoding can substantially alter feature importance rankings. Despite these limitations, SHAP remains one of the most widely used explainability methods and was applied in this project to analyse feature importance.

A specialised variant of SHAP, called TreeSHAP, was also employed. Tree-SHAP was developed specifically for decision-tree-based models and ensembles such as Random Forest and XGBoost. Lundberg et al. (2020) showed that many existing tree-specific attribution methods are inconsistent and proposed TreeSHAP as the only consistent and locally accurate feature attribution method within a broad class of additive explanations. Furthermore, TreeSHAP enables the exact computation of SHAP values for tree ensembles in polynomial time, providing substantial improvements in computational efficiency while retaining the theoretical guarantees of Shapley values (Lundberg et al., 2020). For these reasons, TreeSHAP is particularly well suited to explaining tree-based machine learning models.

The final method was LIME (Local Interpretable Model Agnostic Explanations), which generates local explanations for individual predictions by approximating the behaviour of a complex machine learning model using a simpler interpretable model around a specific instance (Ribeiro, Singh and Guestrin, 2016). Unlike SHAP, TreeSHAP and MDI, which can be used to derive global feature importance rankings, LIME focuses primarily on explaining why a single instance (a compound, or drug, in this work) was classified in a particular way.

## 3. Materials and Methods

### 3.1 The DrugAge Database

The primary source of lifespan-related compound data used in this project was the DrugAge database, which is a curated repository of compounds experimentally associated with lifespan modulation in model organisms (Barardo et al., 2017; Tacutu et al., 2018). The database includes information regarding compounds that either increase or decrease lifespan across multiple organisms including *C. elegans* (worm), fly, mouse, and yeast. For this study, only compounds associated with *C. elegans* were considered in order to maintain biological consistency across the dataset and reduce variability introduced by combining multiple organisms with different biological characteristics. *C. elegans* is widely used in ageing research due to its short lifespan, well characterised genetics, and conserved ageing pathways, making it a valuable model for computational studies of lifespan modulation (Ribeiro et al., 2023). In addition, *C. elegans* is also the model organism for which there is the largest amount of data in DrugAge.

Although DrugAge provided the initial source of lifespan-related compounds, the final dataset used throughout this project was based on a pre-processed dataset provided during project supervision. This dataset already contained binary class labels indicating whether compounds extended the lifespan of *C. elegans* and additionally included verified STITCH identifiers for each compound. These STITCH identifiers were then used to retrieve the values of compound-protein interactions probabilities from the STITCH database (see Section 3.2), and those values were used to create the features in the dataset used in the experiments.

The supervised dataset consisted of 1,120 compounds, which is substantially larger than the 700 lifespan-extension compounds directly obtainable from DrugAge alone. This larger dataset size is consistent with the dataset reported by Ribeiro et al. (2023), since additional negative class compounds were incorporated from the ageing literature in order to provide biologically meaningful non-lifespan-extending examples (instances) and for the models to learn discriminative predictive patterns associated with lifespan extension. Therefore, using experimentally validated compounds allowed the machine learning methods to learn biologically meaningful lifespan-related patterns derived from real experimental evidence.

### 3.2 The STITCH Database

The protein–compound interaction information used throughout this project was obtained from the STITCH database, which integrates protein–chemical interaction evidence from multiple sources. (Kuhn et al., 2012; Szklarczyk et al., 2016). STITCH is a large-scale biological resource that integrates information on chemical–protein interactions from multiple evidence sources, including experimental data, pathway databases, text mining, and computational predictions. For each compound, STITCH provides confidence scores representing the probability of interaction with a specific protein. These scores range between 0 and 1, where larger values indicate stronger confidence in the interaction. Within this project, these probabilities were used as predictive features for the machine learning models. STITCH was selected because it provides bio-logically interpretable compound–protein interaction data and large numbers of protein features suitable for machine learning analysis (Ribeiro et al., 2023; Szklarczyk et al., 2016). The use of protein interaction features also enabled the explainability methods applied in this project to identify proteins that contributed most strongly to model predictions, supporting interpretation of the learned classification models.

### 3.3 Dataset Construction

The final dataset used in this project was constructed by combining lifespan-related compound information obtained from DrugAge (for creating the class labels in the dataset) with protein-compound interaction probabilities obtained from STITCH (for creating the features in the dataset). Each row within the dataset represented a single compound, while each column represented a protein-interaction feature derived from STITCH confidence scores. Consequently, the machine learning models attempted to identify whether specific protein-interaction patterns were associated with lifespan extending compounds.

Before machine learning training, an additional preprocessing and feature filtering stage was applied in order to remove protein-interaction features with low statistical support. More specifically, all protein-interaction features for which fewer than 10 compounds had an interaction probability greater than or equal to 0.5 were removed from the dataset. The motivation behind this filtering procedure was that, after binarisation using a threshold of 0.5, such features would contain too few positive instances to provide reliable statistical support for machine learning generalisation. Consequently, these low statistical support features were considered unlikely to contribute robust predictive information and might instead introduce noise into the classification models.

Following this filtering stage, the number of protein-interaction features was reduced from originally 9,920 to 2,389 features (i.e., the filtering process removed 7,531 protein features). This preprocessing stage therefore reduced feature sparsity while retaining proteins with stronger statistical support across the dataset.

The final dataset consisted of 1,120 compounds represented using 2,389 protein-interaction features. The class distribution was moderately imbalanced, consisting of 867 non-lifespan extending compounds (Class 0) accounting for 77.4% of the instances and 253 lifespan-extending compounds (Class 1) accounting for 22.6% of the instances. This class distribution was highly consistent with the dataset reported by Ribeiro et al. (2023). Additionally, the moderate imbalance between positive and negative classes motivated the use of evaluation metrics such as AUROC and AUPRC, which are generally more informative than classification accuracy alone for imbalanced biological datasets.

Since the dataset already consisted of normalised probability values between 0 and 1, additional feature scaling was not required for the decision tree-based machine learning models used throughout this study. Throughout all experiments, stratified cross-validation was applied in order to preserve class distributions across training and testing folds. Overall, the resulting dataset provided a biologically interpretable representation of compounds through protein-interaction patterns, enabling both predictive modelling and downstream explainability analysis.

### 3.4 Creating Datasets with Different Feature Representations

To evaluate how different representations of compound-protein interaction in-formation affect predictive performance and explainability, three alternative feature representations of the same compound–protein interaction dataset were investigated. All representations contained the same compounds and class labels but differed in how interaction probabilities were encoded.

#### 3.4.1 Continuous Probability Representation

This representation used the original STITCH interaction probabilities directly as features, retaining each confidence value between 0 and 1. While this preserves biological information and allows algorithms to exploit variation in interaction strength, it may also include weak or noisy signals, motivating the evaluation of binary representations.

#### 3.4.2 Binary Fixed Threshold Representation

Continuous probabilities were transformed into binary variables using a fixed threshold of 0.5: values *≥* 0.5 became 1 and all others became 0. This binarisation can reduce noise from weak interactions, allowing models to focus on biologically confident compound–protein associations while simplifying the feature space.

#### 3.4.3 Binary Optimised Threshold Representation

This representation binarised interaction probabilities using an adaptive thresh-old selected during cross-validation. Candidate thresholds of 0.50, 0.55, 0.60, 0.65 and 0.70 were evaluated using only the training data. Within each fold, the threshold achieving the highest validation accuracy was selected and subsequently applied to the full training and test data. The test fold was not used during threshold selection, preventing information leakage and ensuring unbiased performance estimates. A pseudocode description of this procedure is provided in Figure 1 in the Appendix. The aim was to determine whether dataset-specific threshold optimisation could improve predictive performance compared with a fixed threshold of 0.50.

### 3.5 Classification Models

Two machine learning algorithms were investigated in this study: Random Forest and XGBoost. As discussed in Section 2.2, Random Forest is a bagging-based ensemble learning method, whereas XGBoost is a boosting based ensemble learning method.

Random Forest models were implemented using the Scikit-learn library (Pedregosa et al., 2011). The number of trees was set to 500 in order to improve predictive stability and reduce variance. The random seed was fixed at 42, while all remaining hyperparameters were left at their default values. However, XGBoost models were implemented using the XGBoost library (Chen and Guestrin, 2016). The number of trees was also set to 500 to provide stable model estimates. In addition, a learning rate of 0.05, maximum tree depth of 6, subsampling rate of 0.8, and column sampling rate of 0.8 were used to improve generalisation performance and reduce the risk of overfitting. The random seed was fixed at 42, while all remaining hyperparameters were left at their default values.

Model performance was evaluated using stratified 10-fold cross-validation. This ensured that the proportion of positive and negative lifespan-extending compounds remained approximately consistent across all folds. During each iteration, nine folds were used for training and one-fold was used for testing, such that every compound appeared exactly once within a test set. This procedure reduced the risk of overestimating predictive performance and provided a more reliable estimate of model generalisation ability.

The inclusion of both Random Forest and XGBoost enabled comparison between bagging based and boosting based ensemble learning approaches. Furthermore, comparing the behaviour of these algorithms across different feature representations helped evaluate how feature representation affects both predictive performance and the identification of important proteins.

### 3.6 Evaluation Metrics

Several evaluation metrics were used in order to assess predictive performance comprehensively across all classification models. Classification accuracy was used to measure the proportion of correctly classified compounds across the test datasets. Accuracy is a simple overall performance measure, but it may be misleading when datasets contain class imbalance (McDermott et al., 2024).

To address this limitation, the Area Under the Receiver Operating Characteristic Curve (AUROC) was additionally used. This is because AUROC evaluates the ability of a model to discriminate between positive and negative classes across different classification thresholds. A larger AUROC values indicate stronger discriminatory performance. The Area Under the Precision Recall Curve (AUPRC) was also used because it is particularly informative for imbalanced biological classification problems. Precision-recall analysis focuses specifically on the performance of the prediction of positive class, which in this project corresponds to lifespan extending compounds. Since identifying positive lifespan compounds is biologically more important than correctly classifying all negative compounds, AUPRC provides an important complementary evaluation measure (Saito and Rehmsmeier, 2015). All evaluation metrics were averaged across the 10 test sets of the 10 cross-validation folds in order to provide robust estimates of predictive performance and model generalisation ability.

### 3.7 Model Explainability Pipeline

A major focus of this project was not only to achieve strong predictive performance, but also to identify the features (proteins) contributing most strongly to lifespan-extension predictions. Consequently, four feature importance methods were applied throughout the experiments:

Mean Decrease in Impurity (MDI), SHAP, TreeSHAP, and Global LIME. The theoretical foundations of these methods were reviewed in Section 2.3; this section focuses on their implementation within the experimental pipeline. MDI was obtained directly from the feature importance scores generated by the Random Forest and XGBoost implementations available in the Python Scikit-learn library and XGBoost library, respectively. For each trained model, features were ranked according to their importance scores, and the top-ranked proteins were recorded for subsequent analysis. SHAP and TreeSHAP explanations were generated using the Python SHAP library developed by Lundberg and colleagues, Lundberg, Lee et al. (2023). Since both Random Forest and XGBoost are tree-based ensemble models, TreeSHAP was used to efficiently compute feature attribution values for individual predictions. Global feature importance rankings were then obtained by calculating the mean absolute SHAP value of each protein across all instances. Finally, Global LIME explanations were generated by computing local LIME explanations for every compound appearing within the test folds across cross validation (Ribeiro, Singh and Guestrin, 2016). Since each compound appeared exactly once within a test fold, this allowed explanations to be generated for the entire dataset. The local protein importance values generated for individual compounds were subsequently averaged across all instances in the entire dataset in order to produce global LIME based protein (feature) importance rankings. The use of multiple explainability methods allowed comparison of feature importance consistency across the three types of feature representations investigated in this work (as discussed earlier) and across two types of classification algorithms (Random Forest and XGBoost). Proteins consistently identified as top-ranked features across MDI, SHAP, TreeSHAP, and LIME were considered more biologically reliable lifespan associated candidates (Lundberg and Lee, 2017; Salih et al., 2024). Overall, the experimental design enabled evaluation of predictive performance and explainability across different feature representations.

## 4. Results

This section presents the predictive performance and explainability results obtained from the machine learning experiments performed throughout this project. Random Forest and XGBoost models were evaluated across three different feature representations, consisting of continuous protein interaction probabilities, binary features generated using a fixed threshold of 0.5, and binary features generated using optimised thresholds during cross validation, as described in Section 3.4.

We assessed the classification models’ performance using Classification Accuracy, AUROC, and AUPRC averaged across the test sets of a stratified 10-fold cross validation. Explainability analysis was conducted using Mean Decrease in Impurity (MDI), SHAP, TreeSHAP, and Global LIME to identify proteins consistently associated with lifespan-extending compounds. The results are presented separately for Random Forest and XGBoost models, in subsections 4.1 and 4.2, respectively. Moreover, the AUROC and AUPRC curves corresponding to all Random Forest and XGBoost experiments are provided in Appendix.

### 4.1 Results for Random Forest

Although all three feature representations achieved broadly similar performance, the binary representation generated using a fixed threshold of 0.5 consistently produced the strongest results across all evaluation metrics. This representation achieved the highest AUROC (0.7391), AUPRC (0.5202), and Accuracy (0.7884), outperforming both the continuous and optimised-threshold representations. In contrast, the optimised-threshold representation produced the lowest overall performance. The relatively small differences between representations indicate that Random Forest was capable of learning predictive patterns from both continuous and discretised protein-interaction features. However, the consistent superiority of the fixed-threshold representation suggests that binarisation may provide a modest advantage for Random Forest when predicting lifespan-extending compounds.

The top-ranked proteins (features) identified using Mean Decrease in Impurity (MDI) are shown in Table 2. Several differences were observed between the feature representations. The continuous representation identified proteins associated with metabolic and stress related pathways, including *hif-1*, *age-1*, *gsr-1*, and *pdhk-2*. In contrast, the binary feature representations more frequently identified receptor and signalling related proteins, including *ser-1*, *dop-4*, *ser-7*, and *tyra-3*. Some proteins additionally appeared across multiple feature representations, particularly *F55B11.1*, *cca-1*, and *ser-1*, suggesting that these proteins may represent stable predictive signals associated with lifespan-extension within the Random Forest models.

**Table 1:** Predictive Accuracy Results for different feature types using RF.

| Feature Type | AUROC | AUPRC | Accuracy |
| --- | --- | --- | --- |
| Continuous (probability) | 0.7328 | 0.5132 | 0.7875 |
| Binary (0.5 fixed threshold) | **0.7391** | **0.5202** | **0.7884** |
| Binary (Optimal Threshold) | 0.7242 | 0.4993 | 0.7830 |

**Table 2:**
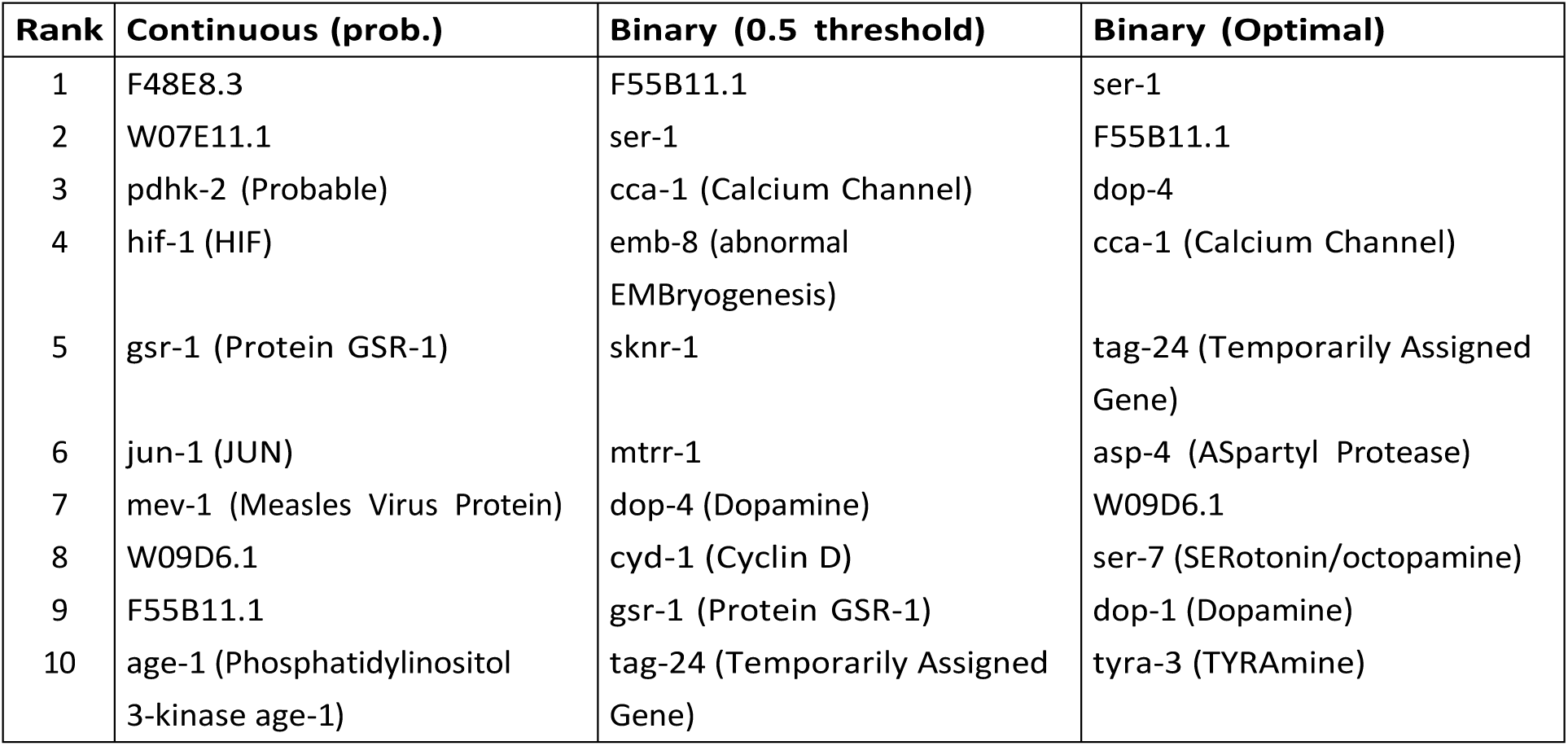
TOP 10-Features (protein names) in the feature ranking produced by Mean Decrease in Impurity (Random Forest)

The protein rankings generated using SHAP and TreeSHAP are shown in Tables 3 and 4, respectively. Substantial agreement was observed between the two explainability methods across all Random Forest feature representations. For the continuous representation, SHAP and TreeSHAP produced identical top-ranked proteins, including *F48E8.3*, *ser-7*, *ser-5*, *pdhk-2*, and *hif-1*. Similarly, strong overlap was observed for the optimised threshold representation, where proteins such as *ser-5*, *cca-1*, *ador-1*, *ser-1*, and *emb-8* consistently appeared across both methods. The binary feature representations more frequently identified receptor and signalling-related proteins, including *ser-1*, *ser-5*, *ser-7*, *ador-1*, and *gnrr-1*, whereas the continuous representation retained a larger number of proteins associated with metabolic and stress-related processes. Overall, the substantial overlap between SHAP and TreeSHAP across all three feature representations indicates highly stable feature attribution behaviour within the Random Forest models. The consistency of these rankings suggests that the most influential proteins identified by the SHAP-based approaches were largely robust to the specific implementation used to compute feature attributions.

**Table 3:**
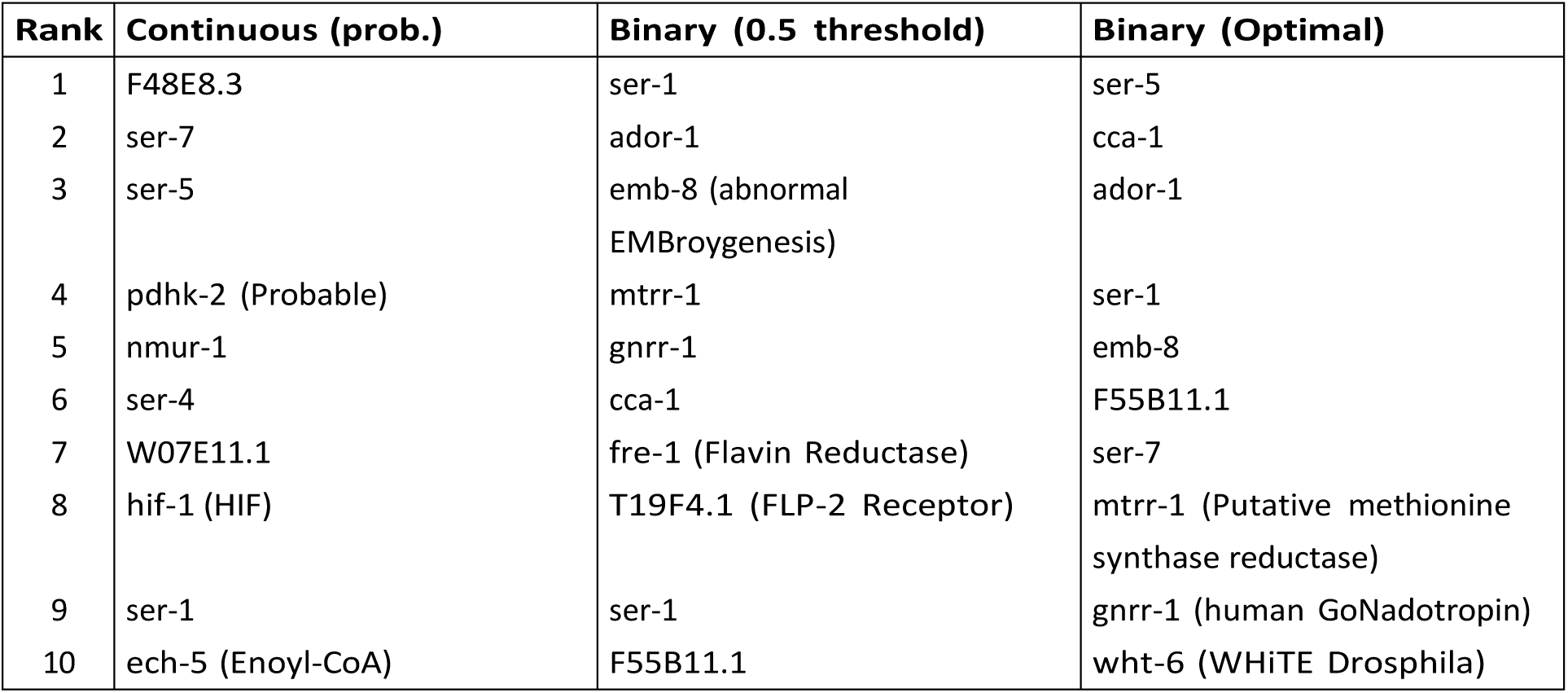
Top 10-Features (protein names) in the feature ranking produced by SHAP (Random Forest)

| Rank | Continuous (prob.) | Binary (0.5 threshold) | Binary (Optimal) |
| --- | --- | --- | --- |
| 1 | F48E8.3 | ser-1 | ser-5 |
| 2 | ser-7 | ador-1 | cca-1 |
| 3 | ser-5 | emb-8 (abnormal<br>EMBroygenesis) | ador-1 |
| 4 | pdhk-2 (Probable) | mtrr-1 | ser-1 |
| 5 | nmur-1 | gnrr-1 | emb-8 |
| 6 | ser-4 | cca-1 | F55B11.1 |
| 7 | W07E11.1 | fre-1 (Flavin Reductase) | ser-7 |
| 8 | hif-1 (HIF) | T19F4.1 (FLP-2 Receptor) | mtrr-1 (Putative methionine<br>synthase reductase) |
| 9 | ser-1 | ser-1 | gnrr-1 (human GoNadotropin) |
| 10 | ech-5 (Enoyl-CoA) | F55B11.1 | wht-6 (WHiTE Drosophila) |

**Table 4:** Top 10-Features (protein names) in the feature ranking produced by tree-SHAP (Random Forest)

| Rank | Continuous (prob.) | Binary (0.5 threshold) | Binary (Optimal) |
| --- | --- | --- | --- |
| 1 | F48E8.3 | cca-1 | ser-5 |
| 2 | ser-7 | F55B11.1 | cca-1 |
| 3 | ser-5 | gnrr-1 | ador-1 |
| 4 | pdhk-2 (Probable) | mtrr-1 | ser-1 |
| 5 | nmur-1 | ador-1 | emb-8 |
| 6 | ser-4 | emb-8 | F55B11.1 |
| 7 | W07E11.1 | F35H10.10 | ser-7 |
| 8 | hif-1 (HIF) | ser-5 | mtrr-1 (Putative methionine synthase reductase) |
| 9 | ser-1 | ckr-2 (CholecystoKinin) | gnrr-1 (human GoNadotropin) |
| 10 | ech-5 (Enoyl-CoA) | ckr-1 (CholecystoKinin) | wht-6 (WHITE Drosophila) |

Unlike SHAP and TreeSHAP, which produced highly similar rankings, LIME identified partially different protein importance patterns across the feature representations. In Table 5 several proteins identified by SHAP-based methods were also identified by LIME, including *pdhk-2*, *emb-8*, *ser-5*, *ser-7*, *gnrr-1*, and *ador-1*. However, LIME additionally identified proteins such as *daf-36*, *daf-4*, and *akt-1*, which were less prominent within the global feature importance methods. Despite these differences, greater overlap between LIME and the SHAP-based methods was observed for the optimised threshold representation than for the continuous representation, suggesting improved consistency between local and global explainability approaches for this feature representation.

**Table 5:** Top 10-Features (protein names) in the feature ranking produced by LIME (Random Forest)

| Rank | Continuous (prob.) | Binary (0.5 threshold) | Binary (Optimal) |
| --- | --- | --- | --- |
| 1 | pdhk-2 | emb-8 | tag-24 |
| 2 | ech-5 | mtr-1 | emb-8 |
| 3 | hsd-1 | gnrr-1 | mtrr-1 |
| 4 | mev-1 | ador-1 | dop-4 |
| 5 | F48E8.3 | ser-5 | ser-5 |
| 6 | WO9D6.1 | ntr-1 | ser-7 |
| 7 | daf-36 | T19F4.1 | gnrr-1 |
| 8 | ZK892.4 | npr-6 | ador-1 |
| 9 | F13D11.4 | nmur-2 | ser-1 |
| 10 | daf-4 | nmur-1 | akt-1 |

To quantitatively evaluate the agreement between explainability methods, Jaccard similarity coefficients were calculated between the top 10 proteins in the feature rankings generated by each pair of feature importance measures across the three feature representations, in the Random Forest models. The resulting Jaccard similarity coefficients are shown in Table 6. The Jaccard similarity coefficient measures the overlap between two sets and is defined as:

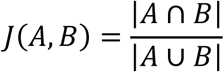

where A and B represent the sets of top 10 proteins identified by two different feature importance methods, |A ∩ B| denotes the number of proteins shared by both sets, and |A ∪ B| denotes the total number of unique proteins across both sets. In this study, the Jaccard coefficient was used to measure how similar the top ranked protein lists were between MDI, SHAP, TreeSHAP, and Global LIME, using the top-10 proteins reported in Tables 2 to 5.

**Table 6:** Jaccard Similarity Coefficients Between Each Pair of Feature Importance Methods across Feature Representations (Random Forest models)

| Explainability Methods | Continuous (models) | Binary (0.5) | Binary (Optimal) |
| --- | --- | --- | --- |
| MDI vs SHAP | 0.2500 | 0.3571 | 0.2500 |
| MDI vs TreeSHAP | 0.2500 | 0.2500 | 0.2500 |
| MDI vs LIME | 0.2500 | 0.0526 | 0.2500 |
| SHAP vs TreeSHAP | 1.0000 | 0.4615 | 1.0000 |
| SHAP vs LIME | 0.1765 | 0.2667 | 0.5385 |
| TreeSHAP vs LIME | 0.1765 | 0.2500 | 0.5385 |

For the continuous probability representation, SHAP and TreeSHAP produced identical sets of top-10 proteins, resulting in a Jaccard similarity coefficient of 1.0. In contrast, much lower overlap was observed between LIME and the SHAP based methods, with SHAP versus LIME and TreeSHAP versus LIME both producing similarity coefficients of 0.1765. The binary representation generated using the fixed threshold of 0.5 demonstrated lower overall agreement between explainability methods. In particular, the overlap between MDI and LIME was extremely low, producing a Jaccard similarity co-efficient of 0.0526. Additionally, SHAP and TreeSHAP agreement decreased substantially compared with the continuous representation, producing a co-efficient of 0.4615. In comparison, the optimised threshold representation produced stronger overall explainability agreement. Similar to the continuous representation, SHAP and TreeSHAP again produced identical sets of top-10 proteins with a similarity coefficient of 1.0. Furthermore, overlap between SHAP based methods and LIME increased substantially, with both SHAP versus LIME and TreeSHAP versus LIME producing coefficients of 0.5385.

Overall, the optimised threshold representation demonstrated the strongest overall agreement between global and local explainability approaches, whereas the fixed threshold representation produced comparatively lower explainability consistency.

### 4.2 Results for XGBoost

Table 7 presents the predictive performance obtained using the XGBoost models across the three feature representations. Overall, good predictive performance was achieved across all experiments, with AUROC values ranging between 0.7299 and 0.7605 and AUPRC values ranging between 0.5056 and 0.5332.

**Table 7:** Predictive Accuracy Results for different feature types using XGBoost.

| Features | AUROC | AUPRC | Accuracy |
| --- | --- | --- | --- |
| Continuous | **0.7605** | **0.5332** | 0.7955 |
| Binary (0.5) | 0.7299 | 0.5164 | 0.8018 |
| Binary (Optimal) | 0.7299 | 0.5056 | **0.8036** |

In contrast to the Random Forest experiments, the continuous probability representation produced the strongest overall predictive performance for XGBoost, achieving an AUROC of 0.7605 and an AUPRC of 0.5332. These findings suggest that preserving the original STITCH interaction probabilities allowed XGBoost to capture more informative biological interaction patterns associated with lifespan extension. Although the binary feature representations achieved slightly higher classification accuracy values, both binary representations produced lower AU-ROC and AUPRC performance compared with the continuous representation. The binary representation generated using the optimised threshold achieved the highest overall accuracy of 0.8036, while the fixed threshold representation achieved an accuracy of 0.8018. However, because AUROC and AUPRC evaluate the ranking ability of the classifier across multiple classification thresholds, the lower values observed for the binary representations suggest that converting interaction probabilities into binary values may have removed useful predictive information from the original STITCH scores. This showed a clear difference between the behaviour of Random Forest and XG-Boost across the feature representations. Whereas Random Forest achieved its strongest predictive performance using the fixed binary threshold representation, XGBoost benefited more strongly from preserving the continuous interaction probabilities. These findings indicate that the effectiveness of a feature representation depends on the underlying learning mechanism of the classifier.

The feature importance rankings generated using Mean Decrease in Impurity are shown in Table 8. Although some overlap existed between the rankings, the continuous and binary representations produced noticeably different sets of highly ranked proteins, indicating that feature representation influenced the proteins identified as important by the XGBoost models. Several proteins appeared consistently across multiple feature representations, including *wah-1* and *dld-1*. The repeated identification of these proteins suggests that they may represent relatively stable predictive signals within the XGBoost models. However, the overall composition of the top-ranked proteins varied considerably across the three feature representations. These results suggest that changes in feature representation affect not only predictive performance but also the interpretation of the resulting models. Overall, the findings indicate that feature representation influences which proteins are emphasised by the model, even when predictive performance remains relatively similar.

**Table 8:** Top 10-Features (protein names) in the feature ranking produced by Mean Decrease in Impurity (XGBoost)

| Rank | Continuous (prob.) | Binary (0.5 threshold) | Binary (Optimal) |
| --- | --- | --- | --- |
| 1 | F48E8.3 | dld-1 | B0334.3 |
| 2 | W07E11.1 | wah-1 | wah-1 |
| 3 | trxr-1 | Y66D12A.9 | C44B7.7 |
| 4 | F20D6.11 | C44B7.7 | dld-1 |
| 5 | F41E6.5 | trxr-1 | C30H6.7 |
| 6 | grk-2 | B0334.3 | R151.10 |
| 7 | gna-1 | ubq-1 | ubq-1 |
| 8 | Y38F1A.6 | dlat-1 | W09D6.1 |
| 9 | C41A3.1 | C30H6.7 (Protein GSR-1) | mocs-1 |
| 10 | F40G9.17 | Y37E3.17 | gsr-1 |

**Table 9:**
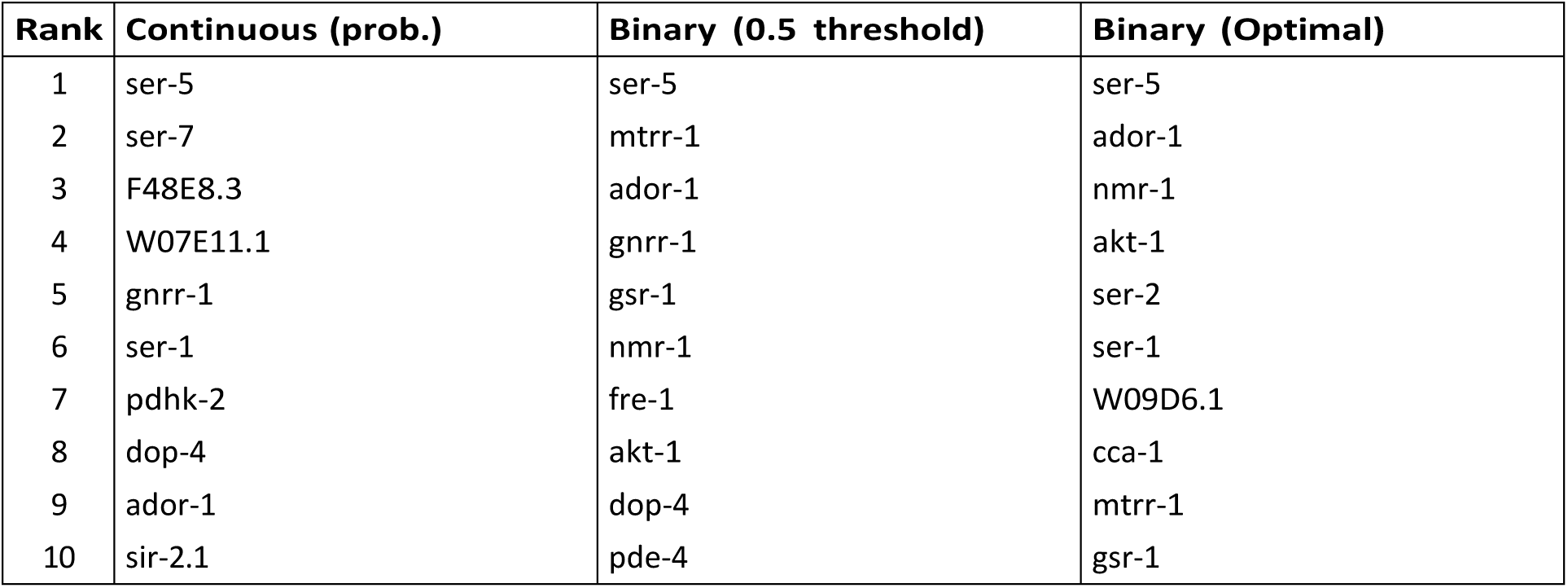
Top 10-Features (protein names) in the feature ranking produced by SHAP (XGBoost)

**Table 10:**
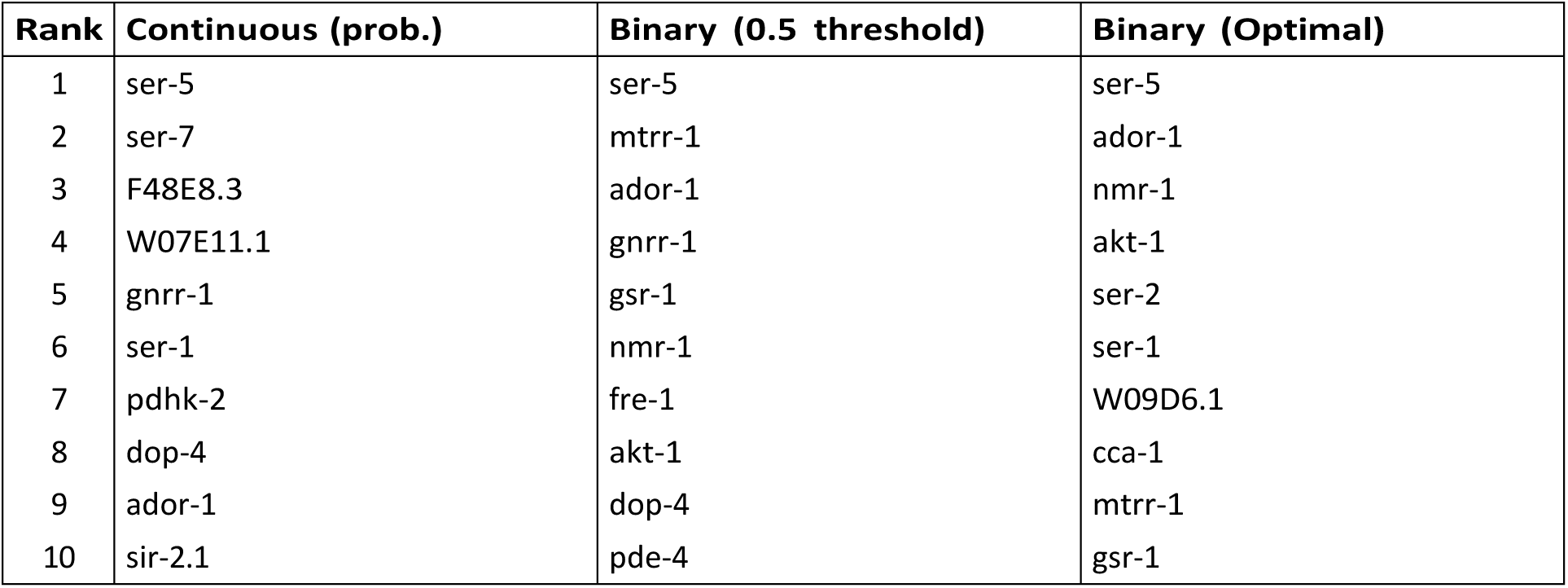
Top 10-Features (protein names) in the feature ranking produced by tree-SHAP (XGBoost)

To further expand the explainability, a particularly notable result was the complete agreement between SHAP and TreeSHAP across all three feature representations. For each representation, both methods produced identical top 10 protein rankings, indicating highly stable feature attribution behaviour within the XGBoost models. This consistency suggests that the most influential features identified by the SHAP based approaches were robust to the choice of explainability method. In addition to the strong agreement between SHAP and TreeSHAP, differences were observed between the continuous and binary feature representations. While several proteins were consistently identified across representations, the overall rankings changed depending on whether the original interaction probabilities were preserved or converted into binary values. This observation supports the first objective of the study, namely to investigate how alternative biological feature representations influence feature importance rankings. A small number of proteins, including *ser-5*, *ser-7*, *ser-1*, and *ador-1*, appeared repeatedly across multiple representations, suggesting that they represent stable predictive signals within the XGBoost models.

The Global LIME rankings shown in Table 11 exhibited some similarities with the SHAP based analyses, particularly through the repeated identification of proteins such as *ser-5*, *ador-1*, *mtrr-1*, *gnrr-1*, and *ser-7* across the binary feature representations. However, LIME also identified several proteins that were not highly ranked by SHAP or TreeSHAP, including *tag-24*, *gar-2*, *dop-1*, and *dop-2*. These differences were also observed across the three feature representations. The continuous representation identified protein such as *pdhk-2*, *F48E8.3*, *ech-5*, and *mev-1*, whereas the binary representations placed greater emphasis on proteins including *ser-5*, *ador-1*, *mtrr-1*, and *gnrr-1* such as *pdhk-2*, *F48E8.3*, *ech-5*, and *mev-1*, whereas the binary representations placed greater emphasis on proteins including *ser-5*, *ador-1*, *mtrr-1*, and *gnrr-1*. These findings further demonstrate that feature representation influences not only predictive performance but also model interpretation and similar patterns were observed in the Random Forest experiments, suggesting that the impact of feature representation extends beyond a specific classification algorithm.

**Table 11:**
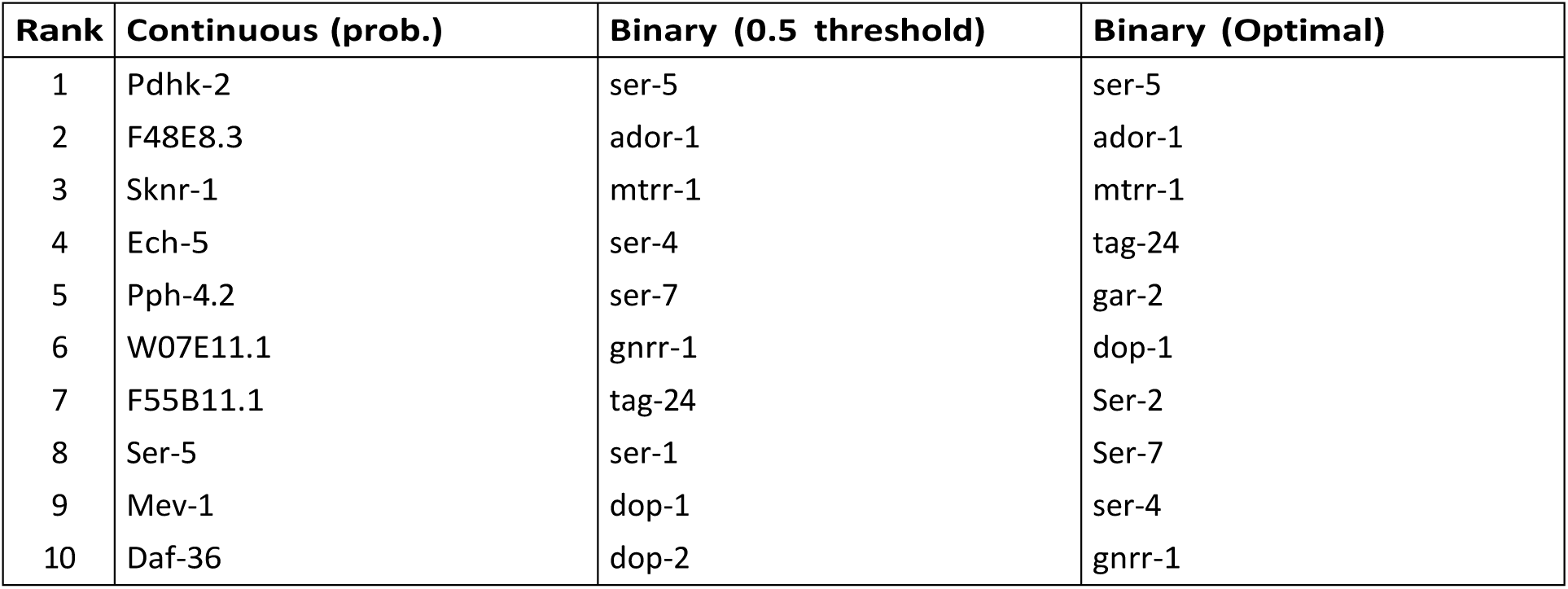
TOP 10-Features (protein names) in the feature ranking produced by LIME (XGBoost)

The continuous probability representation achieved the strongest AUROC and AUPRC performance, whereas the binary representations produced different feature importance rankings and slightly higher classification accuracy values. Taken together, these results indicate that the choice of feature representation affects not only how accurately lifespan extending compounds are predicted, but also how the resulting models are interpreted. Furthermore, the complete agreement observed between SHAP and TreeSHAP provides preliminary evidence regarding the consistency of explainability methods, which is explored further in the subsequent Jaccard similarity analysis.

The Jaccard similarity coefficients calculated between the sets of top-10 proteins identified by the four explainability methods for the XGBoost models are shown in Table 12. Across all three feature representations, SHAP and TreeSHAP achieved perfect agreement, producing a Jaccard similarity coefficient of 1.0. This indicates that both methods identified identical sets of top-ranked proteins irrespective of the chosen feature representation. In contrast, considerably lower agreement was observed between MDI and the SHAP-based approaches. The Jaccard similarity coefficients ranged from 0 to 0.1111, suggesting that MDI frequently identified different proteins from those highlighted by SHAP and TreeSHAP. Moderate agreement was observed between SHAP and LIME, with a Jaccard similarity coefficient of 0.25 across all feature representations. Overall, the Jaccard analysis indicates that explainability behaviour in XGBoost was remarkably stable across feature representations. SHAP and TreeSHAP produced identical sets of top-10 proteins in every experiment, while agreement between SHAP-based methods and LIME remained unchanged across all three feature representations. In contrast to the Random Forest experiments, where explainability agreement varied according to feature representation, the XGBoost models produced highly consistent explanation patterns. This suggests that feature-attribution results derived from boosted tree ensembles may be more robust to changes in feature representation than those derived from bagging-based ensembles.

**Table 12:** Jaccard Similarity Coefficients Between Each Pair of Feature Importance Methods across Feature Representations (XGBoost models)

| Explainability Methods | Continuous | Binary (0.5) | Binary (Optimal) |
| --- | --- | --- | --- |
| MDI vs LIME | 0.1111 | 0 | 0 |
| MDI vs SHAP | 0.1111 | 0 | 0.1111 |
| MDI vs TreeSHAP | 0.1111 | 0 | 0.1111 |
| SHAP vs LIME | 0.25 | 0.25 | 0.25 |
| SHAP vs TreeSHAP | 1 | 1 | 1 |
| TreeSHAP vs LIME | 0.25 | 0.25 | 0.25 |

## 5. Discussion

### 5.1 Effect of Feature Representation on Predictive Performance

The first objective of this study was to investigate how different representations of compound–protein interaction features influence the predictive performance of machine learning models for lifespan extending compound pre-diction. The results showed that feature representation influenced predictive performance, although the magnitude and direction of this effect varied between Random Forest and XGBoost. For the Random Forest models, the binary representation generated using a fixed threshold of 0.5 achieved the strongest overall predictive performance. This representation produced the highest AUROC, AUPRC, and accuracy values among the three Random Forest experiments. One possible explanation is that the binarisation process reduced noise present in the original interaction probabilities, allowing Random Forest to focus on the presence or absence of biologically relevant interactions rather than their exact probability values.

Since Random Forest relies on repeated partitioning of the feature space through decision trees, simplifying the feature representation may have improved the robustness of the resulting ensemble. In contrast, the XGBoost models achieved their strongest AUROC and AUPRC performance using the continuous probability representation.

Unlike Random Forest, XGBoost builds trees sequentially and attempts to correct errors made by previous trees during training. As a result, XG-Boost may be better able to exploit subtle differences in interaction probabilities that are lost when features are converted into binary values. This finding aligns with the theoretical properties of boosting algorithms, which are known to exploit fine grained feature variations through sequential error correction (Friedman, 2001). By contrast, bagging methods such as Random Forest construct independent trees and average their predictions; this approach may benefit from the noise reduction afforded by binary representations, particularly when the original interaction probabilities contain uncertainty or measurement variability. These algorithmic differences underscore that the optimal feature representation is not universal but depends on the learning mechanism employed.

Interestingly, the binary representation generated using the optimised threshold did not outperform the fixed threshold representation in either Random Forest or XGBoost. Although the threshold selection procedure was designed to identify the most suitable binarisation threshold within each cross-validation fold, the resulting models consistently produced slightly lower AUROC and AUPRC values. This may indicate that the threshold optimisation process introduced a degree of overfitting to the training data, reducing generalisation performance on unseen compounds. These findings demonstrate that additional optimisation does not necessarily translate into improved predictive performance. Overall, the results indicate that feature representation is an important design choice in lifespan extending compound prediction. The most appropriate representation depends not only on the biological characteristics of the data but also on the learning algorithm used to construct the predictive model.

### 5.2 Effect of Feature Representation on Explainability

The second objective examined how feature representation influences the identification of important proteins within the learned classification models. Across both Random Forest and XGBoost, the explainability analyses showed that different feature representations produced different sets of highly ranked proteins, indicating that feature representation affects not only predictive performance but also model interpretation. Although several proteins appeared consistently across multiple representations, substantial variation was observed in the composition of the top ranked protein lists generated by MDI, SHAP, TreeSHAP, and LIME. These findings suggest that feature representation influences not only predictive performance but also the biological interpretation of the resulting machine learning models. This observation aligns with recent work demonstrating that SHAP based explanations are sensitive to feature representation, and that simple data engineering operations such as bucketisation or one hot encoding can substantially alter the importance ranks of features (Hwang et al., 2025). In the present study, converting continuous interaction probabilities into binary features similarly changed which proteins were identified as most important, indicating that this sensitivity extends to biological feature spaces. A notable pattern observed throughout the experiments was that continuous representations often retained proteins associated with broader cellular processes, whereas binary representations more frequently emphasised signalling and receptor-related proteins. While detailed biological interpretation is beyond the scope of this study, these observations indicate that the representation of compound–protein interaction data can influence the type of biological information highlighted by explainability methods. Consistent with findings in computational biology, data representation choices influence downstream biological interpretations, not just clustering performance (Weber and Robin-son, 2016). The present study extends this principle to lifespan extending compound prediction, showing that feature representation affects both protein rankings and the agreement between explainability methods. From a machine learning perspective, this result is important because explainability analyses are frequently used to support biological interpretation and hypothesis generation. If feature representation changes the proteins identified as most important, then the conclusions drawn from explainability analyses may depend partly on how the underlying data are represented. Consequently, feature representation should be considered an important component of explainable machine learning pipelines rather than merely a pre-processing step.

### 5.3 Comparison of Explainability Methods

The third objective of this research was to investigate the extent to which different explainability methods produce similar feature rankings when applied to the same classification model. To address this objective, Jaccard similar-ity coefficients were calculated between the sets of top 10 proteins identified by MDI, SHAP, TreeSHAP, and LIME. The strongest and most consistent agreement was observed between SHAP and TreeSHAP. For the XGBoost experiments, SHAP and TreeSHAP produced identical sets of top-10 proteins across all feature representations, resulting in a Jaccard similarity coefficient of 1.0. Similarly, perfect agreement was observed for the continuous and optimised threshold representations in the Random Forest models. These results are expected because TreeSHAP is a specialised implementation of SHAP for tree-based models. (Lundberg, Erion and Lee, 2019) demonstrated that Tree-SHAP computes exact Shapley values for tree ensembles while substantially reducing computational complexity, and (Laberge and Pequignot, 2022) formally explained how and why Interventional TreeSHAP correctly computes these attributions.

The high degree of agreement observed throughout this study therefore provides empirical support for the theoretical relationship between SHAP and TreeSHAP and supports the use of TreeSHAP in tree-based ensembles due to its substantially greater computational efficiency. In contrast, lower levels of agreement were observed between MDI and the SHAP-based methods. This difference is likely to arise because the methods quantify feature importance using fundamentally different principles. MDI evaluates the contribution of features to reductions in class impurity during model construction, whereas SHAP-based methods estimate the contribution of features to individual predictions. Consequently, MDI may favour features that are useful for partitioning the feature space, while SHAP-based methods emphasise features that most strongly influence model predictions. The lower Jaccard coefficients therefore suggest that different explainability methods may capture different aspects of model behaviour rather than identifying a single objective notion of feature importance. Moderate agreement was observed between SHAP-based methods and LIME. While several highly ranked proteins appeared consistently across the methods, notable differences were also observed. This is expected because LIME generates local surrogate explanations around individual predictions, whereas SHAP and TreeSHAP provide feature attributions derived from cooperative game theory. In this study, local LIME explanations were aggregated to produce global protein rankings, which may explain the partial overlap with SHAP-based methods. (Dieber and Kirrane, 2020) noted that the usefulness of LIME explanations for tabular data depends on factors such as feature representation and explanation settings, which may contribute to variation in the resulting feature rankings. Nevertheless, the presence of overlapping proteins across the methods suggests that a subset of proteins consistently contributed to model pre-dictions regardless of the explainability technique employed. An additional finding was that explainability consistency itself was influenced by feature representation. For example, in the Random Forest experiments, the optimised threshold representation produced substantially greater agreement between SHAP-based methods and LIME than the continuous representation. This observation suggests that feature representation affects not only predictive performance and feature rankings but also the degree of agreement between explainability methods. Consequently, explainability results should be interpreted in conjunction with data representation choices rather than independently of them. The variability observed between MDI, SHAP, and LIME, particularly the lower Jaccard coefficients for MDI versus SHAP and SHAP versus LIME, is consistent with prior findings that explainability methods can produce divergent feature attributions under similar conditions (Adebayo et al., 2020). The results presented here indicate that this phenomenon extends to lifespan-extending compound prediction, where different explainability methods may highlight different proteins despite analysing the same underlying model. Furthermore, explainability agreement appeared more stable in XGBoost than in Random Forest, as SHAP and TreeSHAP produced identical sets of top-10 proteins across all XGBoost feature representations, whereas their agreement varied across the Random Forest feature representations. This suggests that the choice of classification algorithm may influence not only predictive performance but also the stability and consistency of model explanations, although further investigation would be required to confirm this observation.

## 6. Conclusion

This work investigated the use of Random Forest and XGBoost models to predict lifespan-extending compounds in C. elegans using compound–protein interaction features derived from the STITCH database (Szklarczyk et al., 2016). Three alternative feature representations were evaluated, and model interpretation was analysed using four explainability methods: MDI, SHAP, TreeSHAP, and LIME. The results demonstrated that feature representation influences both predictive performance and model interpretation.

For Random Forest, the binary representation generated using a fixed threshold of 0.5 achieved the strongest predictive performance, whereas XGBoost achieved its strongest AUROC and AUPRC performance using the continuous probability representation. These findings indicate that the effectiveness of a feature representation depends on the underlying machine learning algorithm, which is consistent with the differing learning mechanisms of bagging and boosting approaches (Friedman, 2001).

The explainability analyses showed that different feature representations produced different sets of highly ranked proteins across both Random Forest and XGBoost models. This demonstrates that feature representation affects not only predictive performance but also the interpretation of machine learning models. These findings are consistent with recent work showing that explainability results can be sensitive to feature representation and feature engineering choices (Hwang et al., 2025), and with previous observations that data representation influences downstream biological interpretation (Weber and Robinson, 2016). The comparison of explainability methods showed that SHAP and TreeSHAP produced the most consistent results across feature representations and classification algorithms. This finding is supported by the theoretical properties of TreeSHAP, which computes exact Shapley values for tree-based models while maintaining the guarantees of the SHAP framework (Lundberg, Erion and Lee, 2019; Laberge and Pequignot, 2022).

Lower levels of agreement were observed between MDI, SHAP, and LIME, supporting previous findings that different explainability methods can produce substantially different feature importance results (Adebayo et al., 2020; Dieber and Kirrane, 2020). Overall, this study demonstrates that both feature representation and explainability method selection can influence the conclusions drawn from machine learning models in biological applications. The findings contribute to a better understanding of how compound–protein interaction data can be represented for lifespan-extending compound pre-diction and provide insights into the consistency and reliability of different explainability approaches.

## 7. Limitations and Future Work

Several limitations should be considered when interpreting the findings of this study. First, the experiments were conducted exclusively using data from C. elegans. Although this organism is widely used in ageing research, the extent to which the findings generalise to other organisms remains uncertain. Recent studies have highlighted the importance of organism-specific biological mechanisms when applying machine learning to ageing research (Ezcurra et al., 2024). Secondly, the study relied solely on compound-protein interaction features derived from the STITCH database (Kuhn et al., 2012; Szklarczyk et al., 2016). While STITCH provides a large and biologically meaningful set of interaction features, it does not capture other potentially relevant sources of information, such as gene expression data, pathway information, or other omics data.

Additionally, although multiple explainability methods were evaluated, the biological significance of the identified proteins was not experimentally validated. Consequently, the feature importance rankings should be interpreted as evidence of model behaviour rather than definitive confirmation of biological importance. This is consistent with broader reviews of AI applications in ageing research, which highlight the need for experimental validation of computational findings (Mahbub, Safaeian and Sohrabi, 2026). Finally, model performance was evaluated using stratified 10-fold cross-validation rather than independent external datasets, which would provide stronger evidence regarding the generalisability of the proposed approaches.

Future research could integrate additional biological data (e.g., pathways, Gene Ontology, gene expression) to improve lifespan prediction (Ezcurra et al., 2024). More advanced machine learning, such as graph neural networks, may better capture compound protein interactions. A wider range of explainability methods and similarity measures could also be explored, alongside rigorous benchmarking and biological validation (Mahbub, Safaeian and Sohrabi, 2026).

Beyond these directions, this study has identified a specific testable hypothesis: the protein ser 5 (serotonin receptor) was top ranked across all XGBoost models and most Random Forest representations. A recent preprint (Avchaciov et al., 2024) showed that lifespan extension by Zotepine and Ziprasidone depends partially on ser 5 in C. elegans, confirming its biological relevance. However, no study has used machine learning to rank ser 5 as a predictive feature or proposed specific, untested compounds interacting with ser 5 based on interaction probabilities. Thus, our approach independently identifies ser 5 as a robust signal and generates novel predictions: compounds from our best model (continuous XGBoost, AUROC 0.7605) that strongly interact with ser 5 are high-priority candidates for an in vivo validation. Finally, experimental validation of top ranked proteins and predicted compounds would provide crucial biological confirmation. These directions can enhance predictive performance, interpretability, and the application of explainable ML in ageing research.

## Appendix

This Appendix consists of three figures. Figure A.1 shows the pseudocode for binary-optimised threshold selection. Figures A.2 and A.3 show the effects of different feature representations on AUROC values and AUPRC values, respectively.

**Figure A.1:**
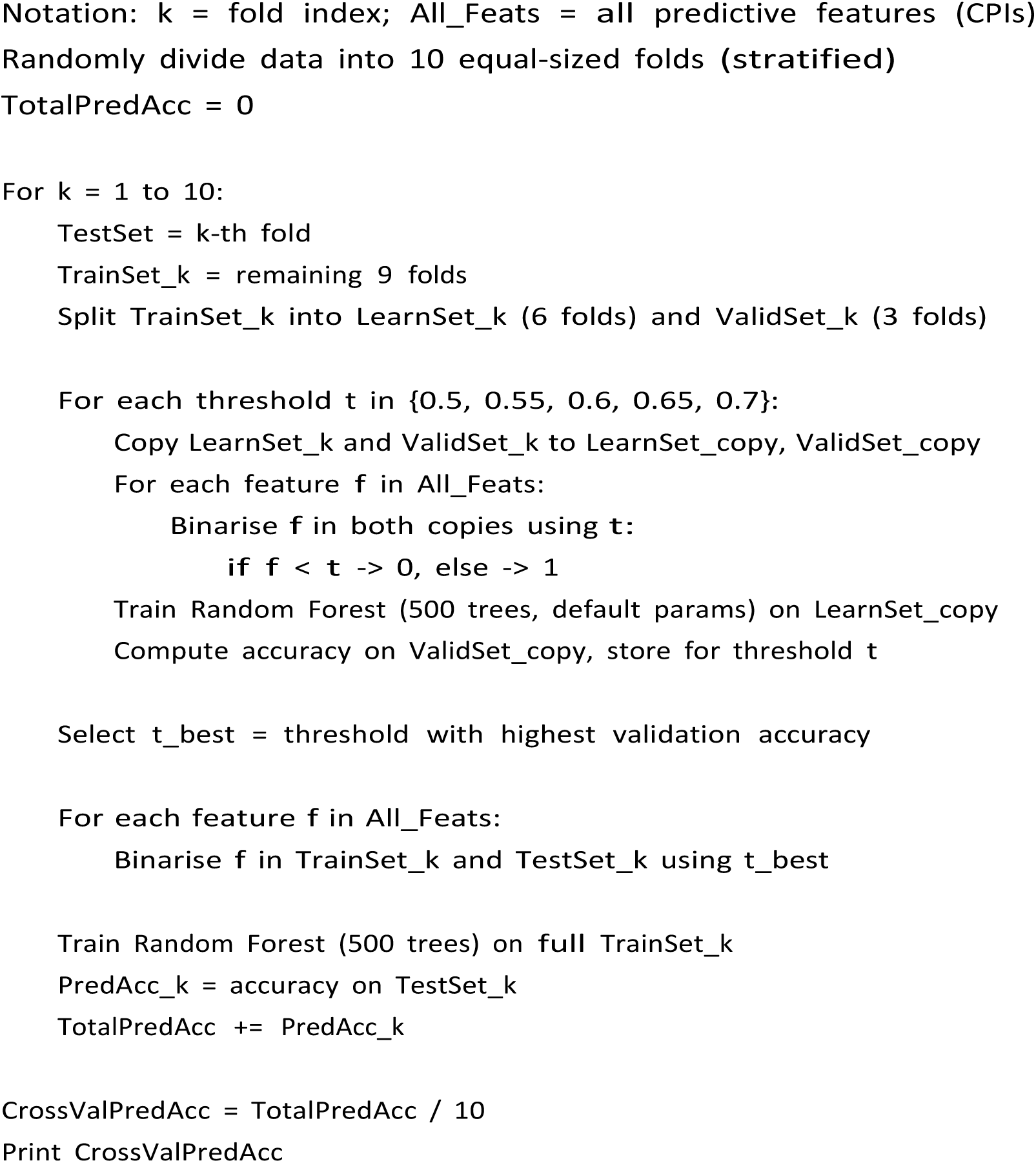
Pseudocode for binary optimised threshold selection using 10-fold cross-validation (applied separately for each fold). This procedure generated the third feature representation described in Section 3.4.3.

**Figure A.2:**
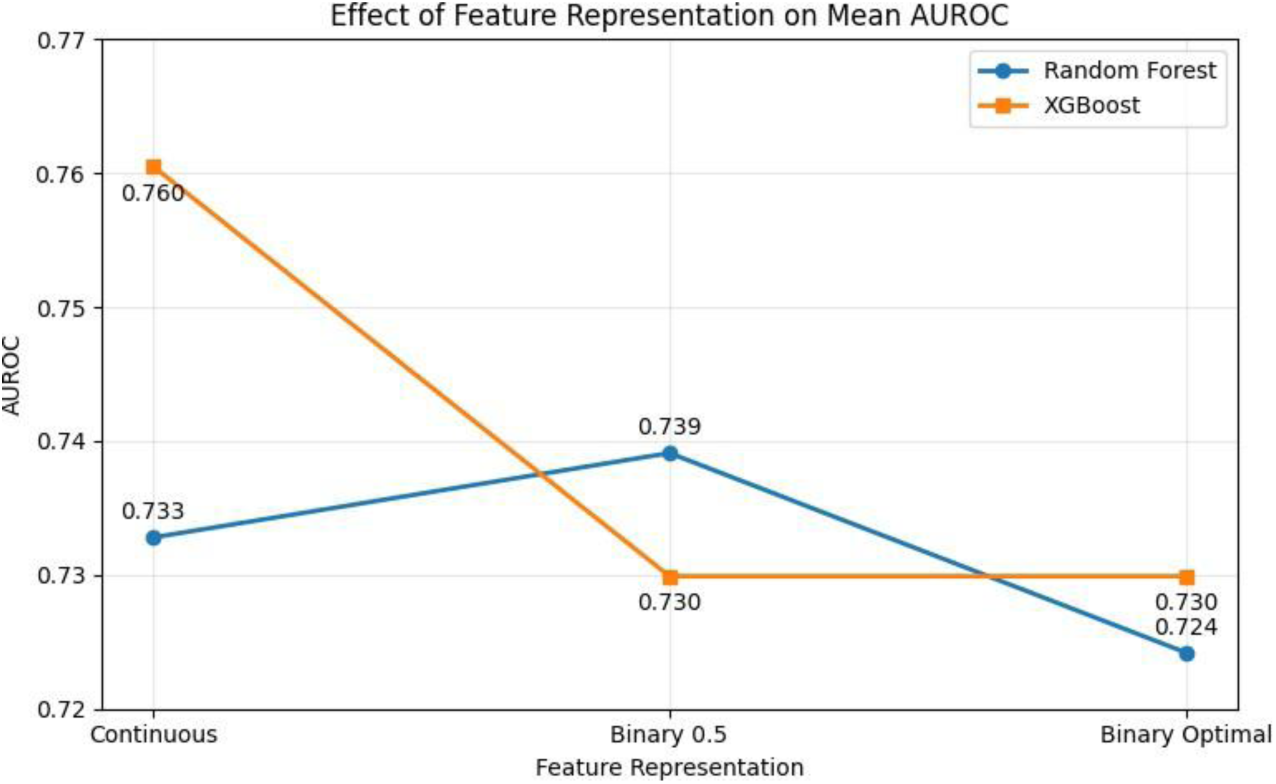
Effect of feature representation on mean AUROC for Random Forest and XGBoost models.

**Figure A.3:**
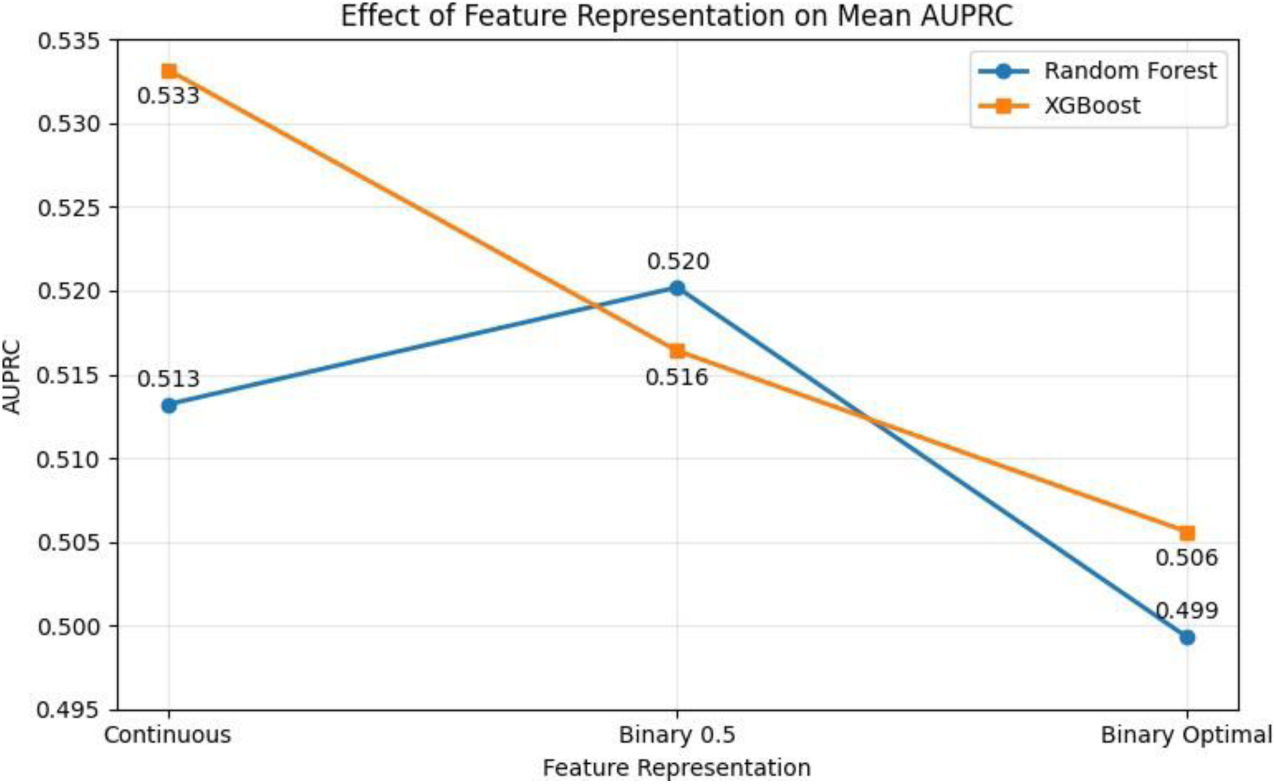
Effect of feature representation on mean AUPRC for Random Forest and XGBoost models

